# Resource-Driven Encounters and the Induction of Disease Among Consumers

**DOI:** 10.1101/091850

**Authors:** Rebecca K. Borchering, Steve E. Bellan, Jason M. Flynn, Juliet R.C. Pulliam, Scott A. McKinley

## Abstract

*Submitted Manuscript 2016.* Territorial animals share a variety of common resources, which can be a major driver of conspecific encounter rates. We examine how changes in resource availability influence the rate of encounters among individuals in a consumer population by implementing a spatially explicit model for resource visitation behavior by consumers. Using data from 2009 and 2010 in Etosha National Park, we verify our model's prediction that there is a saturation effect in the expected number of jackals that visit a given carcass site as carcasses become abundant. However, this does not directly imply that the overall resource-driven encounter rate among jackals decreases. This is because the increase in available carcasses is accompanied by an increase in the number of jackals that detect and potentially visit carcasses. Using simulations and mathematical analysis of our consumer-resource interaction model, we characterize key features of the relationship between resource-driven encounter rate and model parameters. These results are used to investigate a standing hypothesis that the outbreak of a fatal disease among zebras can potentially lead to an outbreak of an entirely different disease in the jackal population, a process we refer to as indirect induction of disease.

## 1 Introduction

Due to the rapid growth in high-resolution animal movement data, there is a growing recognition that classical models for encounter rates among animals should be revisited [1, 2]. Some recent progress on this point has been made in foraging theory. Theoreticians have shown that predictions for search efficiency, and its dependence on prey density, can be substantially different when movement models include intermittent long-range relocation events [3] or local sensing and decision-making [4]. The resulting nonlinear dependence on prey density yields novel forms for functional response in predator-prey systems, and raises questions about what other theoretical frameworks would benefit from the inclusion of more realistic animal behavior.

Understanding the impact of sudden environmental changes on animal behavior is a particularly compelling application. Intentionally or unintentionally, humans frequently alter the availability of resources for consumer species, invariably leading to unintended consequences. As reviewed by Oro et al. [5], considerable work has been devoted to identifying instances of anthropogenic resource provisioning. Other work has examined the impact of naturally occurring resource subsidies that occur as pulses in either space or time on consumer population dynamics, behavior and community structure (Rose & Polis [6], Anderson et al. [7], Clotfelter et al. [8], Holt [9], Ostfeld & Keesing [10], Yang et al. [11]). A smaller but substantial body of work (reviewed by Becker et al. [12] and Sorensen et al. [13]) specifically considers the effects that such provisioning can have for infectious diseases of consumers.

In order to develop a predictive framework for the impact of resource supplementation on the spread of disease among consumers, one must establish a relationship between resource density and a rate of conspecific consumer encounters. This is especially true when considering directly transmitted pathogens such as rabies virus or canine distemper virus. Becker et al. [12] recently considered this relationship explicitly in their review on the link between anthropogenic resources and wildlife-pathogen dynamics. Motivated by the prevailing trend revealed by their meta-analysis, they introduced the assumption that increased resource availability will lead to increased consumer aggregation, which in turn implies increased infection risk when a pathogen is present. In contrast, we highlight that the relationship between resource availability and consumer encounters need not be strictly increasing. At some level of resource availability, consumers may no longer need to share resource sites, meaning that consumer encounters might actually decrease. The question then concerns whether there is a critical resource density above which consumer aggregation no longer increases, and how that density would depend on parameters that can be inferred from data.

With the foregoing discussion as motivation, in this work we investigate the role resources play in helping to maintain pathogen transmission or facilitate disease emergence. Specifically, we quantify the relationship between increases in resource availability and the consequent changes in conspecific encounter rates among consumers. Furthermore, we provide context from disease ecology to address the question “How big is big?” when it comes to encounter rate changes. To this end, we consider the relationship between the potential for disease maintenance among a population of jackals and the annual occurrence of anthrax outbreaks among local herbivores in Etosha National Park (ENP) in Namibia [14,15,16]. This surge in available carcasses serves as a supplemental resource for the local jackal population, and due to two years worth of tracking data, we have new understanding about how jackals respond to temporarily available resources [15]. The jackals live in territorial family groups. They regularly hunt and forage within and nearby their defendable territory, and opportunistically scavenge on carcasses when they are available. In studying the data, we also observe that jackals sometimes make long treks to visit resources (see Figure 1), possibly crossing through the territories of neighboring family groups.

**Figure 1:**
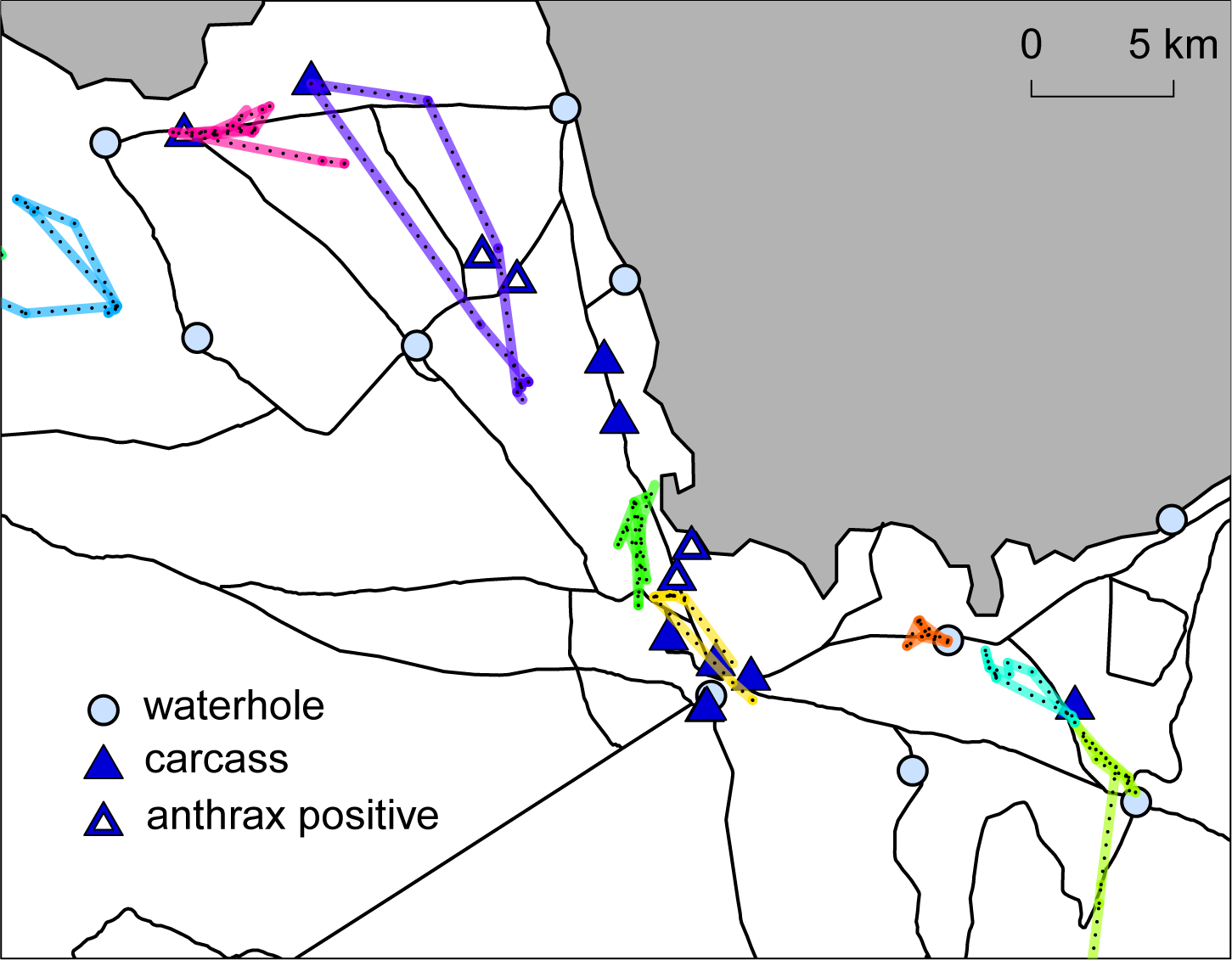
Collared jackal movement and known carcass locations in Etosha National Park. Left: GPS locations for one jackal over one week time frames, beginning on February 2, 2010 (upper) and June 6, 2009 (lower). Right: GPS locations for all collared jackals on February 2, 2010. Jackals are differentiated by color. Each colored line segment with black dashes connects two GPS pings for that jackal. Blue circles represent waterholes. Blue triangles indicate locations of known carcasses. White triangle insets indicate that the carcass tested positive for anthrax causing bacteria. Roads are indicated by black lines and the shaded gray areas are part of a salt pan in ENP.

These movement patterns have interesting implications for the potential spread of disease. It is possible that during resource pulses, jackals will have increased contact with individuals outside their family group. As a consequence, though anthrax bacteria rarely cause disease in carnivores, an intense uptick in jackal-to-jackal encounters could lead to an outbreak of a *different* disease in the jackal population [15]. In this particular sense, we might say that anthrax can “cause” a rabies epidemic. We refer to this process as *indirect induction of disease* because change in resource availability does not introduce a pathogen, it simply changes a population's contact network structure in such a way that the population becomes susceptible to invasion by a pathogen that would not otherwise be able to take hold. In fact, as we argue later, induction of disease can result from a *decrease* in resource availability as well.

## 2 Model development and preliminary analysis

### 2.1 Resource-driven encounters

With the jackal population from ENP in mind, we introduce the following general assumptions:

- the locations of both consumers and resources are randomly distributed throughout a spatial region of interest;
- the resources are only available for a given interval of time *τ*_i_ and new resources are located independently of previous ones;
- consumers are territorial, spending most of their time near a home location, and have a limited range of detection, characterized by a length scale *ℓ*;
- consumers prefer to visit the nearest resource they detect;
- they respond to resources independently of other consumers; and,
- they are satiated after visiting a resource, and therefore visit at most one resource per unit of time *τ*_2_.

We are interested in the number of conspecific encounters a typical consumer will have as a result of temporarily available resources.

For the sake of simplicity, and because we believed the choice was reasonable for the jackal population in ENP, we choose the time parameters to be the same, *τ_1_ = τ_2_ = τ* = one week. We use 𝒪 to denote the spatial region we are studying. For each week, resources are distributed throughout 𝒪 according to a Poisson spatial process, with intensity parameter *κ*. This means that for any region of area *A* contained in 𝒪, the number of resources in that region is Poisson distributed with mean *κ* Moreover, if two regions are disjoint, their respective numbers of resources are independent. We assume there is a consumer located at the origin, referred to as the *focal consumer*. The remaining consumers are distributed throughout 𝒪 according to a Poisson spatial point process with intensity *ρ*. These intensity parameters correspond to the expected population density produced by the model for the consumers and resources, respectively. In our simulation and mathematical analysis, the size of the landscape is taken to be sufficiently large that the presence of a boundary does not have an effect on quantities of interest.

To model the consumer's limited ability to detect resources and/or travel to resources that have been detected, we assume there is a maximum distance *ℓ* within which a given consumer will detect resources. Moreover, we assume that consumers will detect all resources within a surrounding circle of radius *ℓ* and will choose to visit the nearest of these detected resources. To understand the consumer-resource landscape, it is helpful to construct Voronoi tessellations of the region 𝒪 generated using the set of the resource locations [17]. Using the R-package deldir [18], we display three such tessellations in Figure 2. Consumer locations are displayed as squares while resource locations are triangles. The focal consumer appears in black. Each subregion of the tessellation, referred to as a *Voronoi cell*, contains exactly one resource and is comprised of all points that are closer to this local resource than any other. We also refer to a resource's Voronoi cell as its *basin of attraction.* We stress that when resources are rare, the basin of attraction will usually contain many points that are a distance greater than *ℓ* from the resource. If a consumer is located at such a point, it will not visit any resources during that unit interval of time.

**Figure 2:**
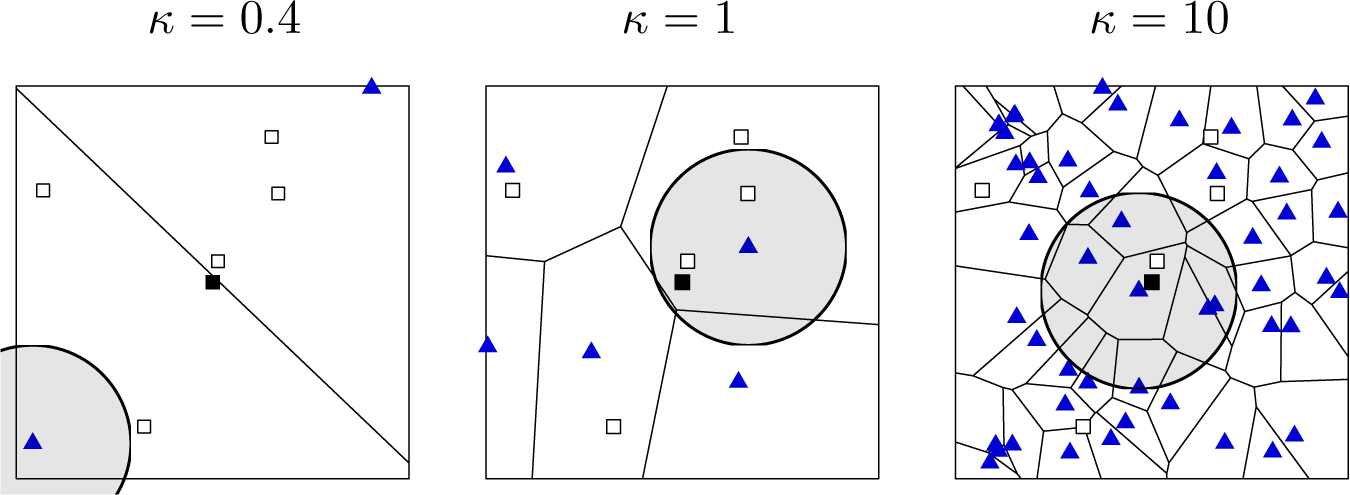
Voronoi diagrams displaying the regions of “attraction” for each resource. Each blue triangle indicates the location of a resource. The black square and the white squares indicate the locations of the focal consumer and non-focal consumers, respectively. Gray circles, centered at the resource closest to the focal consumer, display the region where consumers can detect the resource. From left to right, the number of resources displayed in each panel is 2, 5, and 50.

The fundamental goal in the analysis of our model is to understand the number of encounters that occur due to the presence of a particular type of resource. We define the *resource-driven encounter rate ε* to be the expected number of consumers that choose the same resource as the focal consumer per unit interval of time. In Figure 2, the focal consumer in the left, center and right panels has 2 and 1 encounters respectively. This reveals a fundamental dynamic in the model: that intermediate resource availability can produce the highest encounter rates. When resources are scarce, resource-driven encounters are rare because it is unlikely that the focal consumer is near enough to a resource to detect it. On the other hand, when resources are common, encounters are rare because nearby consumers have local resources of their own to visit.

To estimate ε for a given parameter triplet (*ρ, *κ*, ℓ*), we simulated 1000 independent landscapes, calculated the resulting number of encounters for the focal consumer in each, and then took the average of these observations. For most of our simulations, we used the parameter ranges *ρ ∊* (0,10) and *ℓ ∊* (1,14). As described in Section A.3, for every triplet (*ρ, *κ*, ℓ*) there is an associated triplet (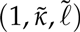) for which *ε* is the same. We therefore always set *ρ* = 1 in our simulations and use the transformation 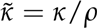 and 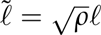 to compute *ε* when *ρ ≠* 1.

### 2.2 Introduction of pathogen and outbreak of disease.

To place our encounter rate results in the context of disease ecology, we employ a simple stochastic model of pathogen spillover between two “adjacent” populations. Because we assume that the disease is initially not present in the target population, we incorporate a rate γ_spinover_ of pathogen introduction events from a maintenance population in which the disease is endemic. Due to our interest in transient seasonal effects, the results are expressed in terms of the duration *Τ* of the resource increase.

We make three central assumptions:

- the time scale of an outbreak is small relative to the time it takes for significant changes in population size to occur;
- each introduction of a pathogen involves just one initial infectious individual; and,
- the arrival times of pathogen spillover events are independent.

Under these assumptions, the initial pathogen invasion process is intrinsically tochastic. We model the introductions as a Poisson arrival process with rate pa-rameter γ_spinover_, an assumption similar to the invasion model proposed by Drury et al. [19]. For the transmission events among individuals in the target population, we use a stochastic Susceptible-Infectious-Susceptible (SIS) model. Because the total population size is fixed in this model, it is only necessary to track the state ransitions for the infectious group, whose population size at time *t* is denoted *I(t)*. The transition rates for our continuous-time Markov chain are given by

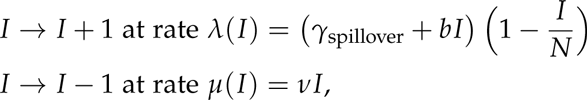

where *N* is the target population size, *v* is the clearance (or disease-related mortality) rate, and *b*(*1 – I/N*) is the average number of transmissions per unit interval of time by an infectious individual when the infectious population has size I.

While this simple model of transmission ignores other potentially relevant characteristics (e.g. latent periods, population turnover, and acquired immunity), our present focus is on how consumer-resource interactions modulate transmission dynamics in the early introduction phase. We are specifically interested in the probability that the level of infection can reach an endemic state in the target population before the period of resource increase dissipates. Given our context that the disease dynamics take place over a large area and the pathogen introductions are relatively rare, we introduce a fourth assumption:

- each pathogen introduction resolves itself independently in the target population (either to extinction or invasion).

Mathematically, this is tantamount to omitting the γ^spinover^ term in the transition rate formulas and treating each pathogen introduction event independently. The “endemic equilibrium” is the minimum size for the infectious population such that the rate of increase equals the rate of decrease. We consider a pathogen introduction to be “successful” if the size of the resulting infectious population eventually exceeds the endemic equilibrium value:

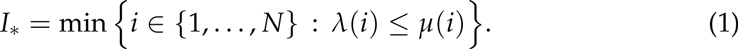

We then study the continuous-time Markov chain *{I(t)}t*_*t*>0_ with the transition rates

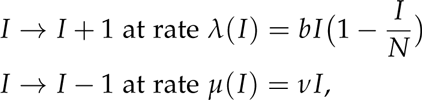

and compute the probability of successful invasion assuming that a pathogen been introduced at time zero:

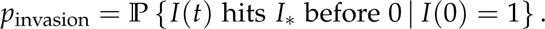

In a sense made rigorous by Kurtz [20], when *N* is large this stochastic system ehaves more and more like an associated ODE,

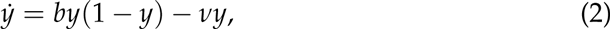

where we interpret *y(t)* as the proportion of the population that is infectious at time *t*. If *b > v* and *y*(0) > 0, then *y(t)* converges to the equilibrium value *y*_*\**_ = 1 *– v/b.* Otherwise *y(t)* → 0 as *t* → ∞. Following the terminology used by Ball [21] (see also Heffernan et al. [22]), we refer to 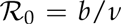 as the *reproductive ratio.*

In contrast to the ODE model, no matter how large *N* is, in the stochastic model there is always a chance that an infectious lineage will go extinct before it reaches an endemic state. In Figure 3 we display ten stochastic SIS paths with a population size of 50 with *b* = 2 and *v* = 1. Some of these paths quickly go extinct, while others reach the endemic state. Overlaid on the stochastic paths is *Ny(t)*, the rescaled solution to the associated ODE (2), with initial condition *Ny(0)* = 1.

**Figure 3:**
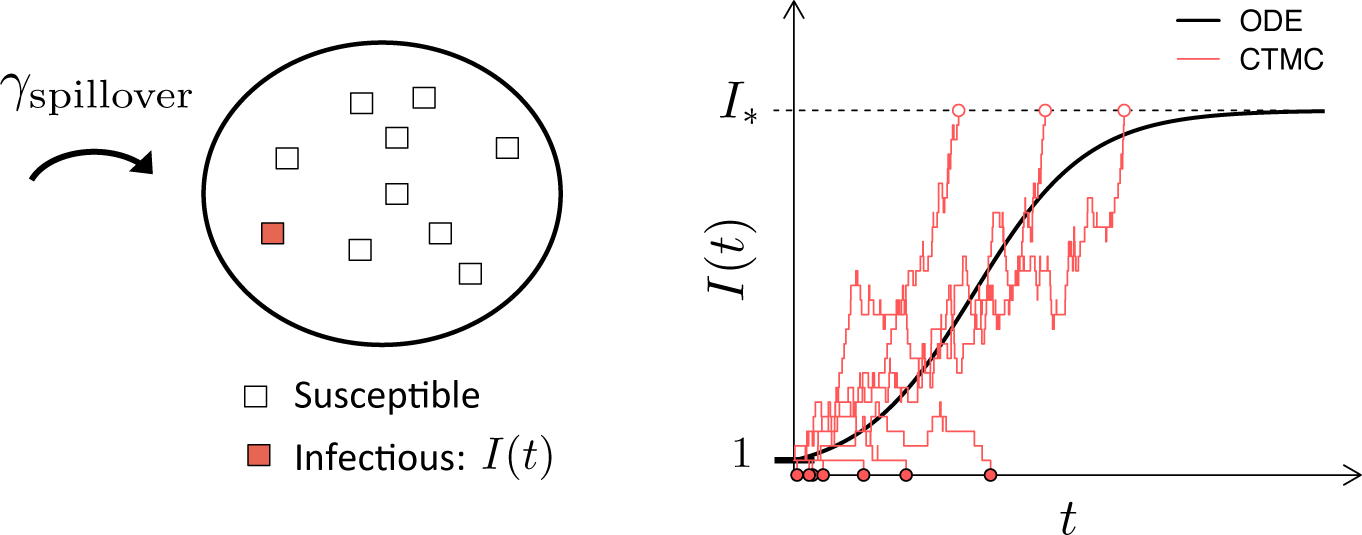
Number of infectious individuals resulting from the introduction of one infectious individual in a population of size *N* = 50. Ten sample paths for the stochastic SIS model defined in Appendix 2.2 are plotted (red lines) with the solution to the analogous ordinary differential equation model (black curve). Our representation for having achieved the endemic state is *I*_*\**_ (dashed horizontal black line), which is defined in Equation 1. Open circles are plotted when *I*_*\**_ is reached before 0 and red points indicate when the pathogen died out of the population before reaching *I*_*\**_.

Just as it is for the ODE model, the reproductive ratio is a critical dimensionless parameter in the stochastic model. When 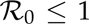, then as *N → ∞*, ρ_invasion_ → 0 [23]. On the other hand, when 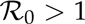, then as *N → ∞*, the probability of invasion is strictly greater than zero. As described in Appendix A.4, ρ_invasion_ is commonly approximated by computing the complement of an extinction probability for an associated branching process [24]. This gives the approximation

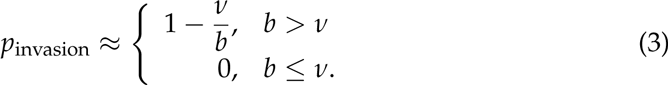

With this in hand, we can estimate the probability that there is a successful pathogen invasion in the target population during the period of increased resource availability, *t ℓ* [0, *T*]. For each pathogen introduction, we label it “successful” with probability ρ_invasion_ and then note that, from Markov chain theory [25] the arrival of *successful* introductions is also a Poisson process, but with a “thinned” intensity γ_spmover_ρ_invasion_. This implies that the time of the arrival of the first successful spillover has an exponential distribution with rate parameter γ_spmover_ρ_invasion_. Therefore, the probability of a successful invasion occurring during the resource pulse has the form

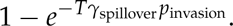

### 2.3 Data collection and analysis

Jackals were captured from January 2009 to July 2010 in central ENP as part of a larger study on jackal movement and anthrax ecology (Bellan et al [15]). Twenty-two adult jackals were fitted with GPS (global positioning system; African Wildlife Tracking, Pretoria, Republic of South Africa) collars based on the requirement that they were large enough to limit the collar to less than 6% of body weight. Movement data was acquired from collars by VHF radio-tracking animals and downloading recorded hourly GPS fixes with UHF download. Due to challenges associated with acquiring downloads, there is some variation in the time intervals between recorded locations. In some cases there are missing data points; and in a few cases, observations were made more frequently than once per hour. The duration of time each collared animal was observed also varied greatly, from a few weeks to 2 years, for a total of 13.5 jackal-years of (roughly) hourly location data.

In addition to jackal position data, carcass surveillance data was recorded from January 2009 to November 2010 (see [15] and [26] for additional information). Multiple characteristics of a carcass were recorded, such as: species, date of observation, level of degradation, and cause of death. Particularly relevant to our investigation, there are jackal counts recorded for 299 out of 411 carcass sites (178 out of 244 zebra carcass sites). These data are displayed in Figure 6.

*Resource visits.* For each recorded instance of a carcass, we assigned a “carcass active interval” based on its estimated time of death and the level of degradation at the time of discovery (if recorded). This window lasted up to six days. Six days also served as the baseline duration of availability, used when low or no level of degradation was recorded. For each jackal that was tracked in the park contemporaneously with a known carcass, we computed a “time-local average position,” i.e. the mean of all recorded positions of the jackal during the carcass active interval. The distance between this average position and the location of the active carcass was assigned to be the distance of a resource visit or non-visit. If the jackal's minimum distance from the carcass of interest during this period was less than 100m, we classified the event as a resource visit.

Our use of the location datasets for collared jackals to identify “resource visits” assumes that visits to carcass sites are meaningfully captured within the movement data. If jackal movement were not influenced by the distribution of carcasses on the landscape, we would expect to find no association between jackal locations and identified carcass sites. We performed a randomization test to assess the null hypothesis that there is no association between jackal locations and identified carcass sites. For each site, we held the time that the site was available constant and reassigned the location by sampling (without replacement) from the recorded carcass locations. For each permuted dataset, we then calculated the number of “resource visits” in the same way as described above for the observed data. Using the true carcass location data there were 10 and 44 visits from jackals with a time-local average of 10+ km and 5+ km respectively from the carcass locations. Out of 1000 sets of randomized carcass locations, the maximum number of 10+ km and 5+ km visits was 6 and 15, corresponding to a *p*-value of < 0.001 for both distances, and indicating that there is a highly significant association between jackal locations and identified carcass sites.

## 3 Results

By way of simulation and analysis, we are able to characterize the most prominent qualitative features of the expected number of encounters experienced by a focal consumer, as it depends on the expected resource density ρ and the maximum distance of detection *𝓁*. While we report the results of the specific model described in the previous section, as long as a model is consistent with the listed assumptions, then our fundamental conclusions are the same: there is a non-monotonic relationship between the expected resource-driven encounter rate and the resource density; the maximum potential encounter rate can be large in terms of its impact on the critical disease ecology parameter 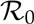; and, somewhat surprisingly, low resource densities are associated with the largest increases in encounter rates.

### 3.1 Analysis of the consumer-resource model

We summarize the predictions of the mathematical model as follows. For fixed values of *ρ* and *𝓁*:

- *From the point of view of an available resource (carcass), the number of visitors decreases with *κ*.* The presence of additional resources increases the number of options for consumers and so, as ρ increases, the expected number of visitors at a given resource site decreases.
- *From the point of view of an individual consumer, the number of encounters increases, then decreases with *κ**. When resources are scarce, most consumers will not be near enough to detect them. Increasing ρ means that more and more consumers visit resources, leading to increased consumer-consumer interactions. The effect is not monotonic though. When resources are abundant, consumers will generally detect more of them. Due to this increase in available options, it becomes less likely that multiple consumers will visit the same resource.

From the point of view of a given resource site, there are two limiting factors on the number of visitors: 1) the size of the resource's basin of attraction, as defined by the Voronoi tesselation described in (Section 2.1 and presented in Figure 2; and 2) the consumers' limited distance of detection. As *κ* increases, the resource's basin of attraction decreases in size, therefore limiting the pool of consumers that would choose it. In Section 3.2 we present an analysis of the ENP data set, wherein we find some evidence that the number of jackals expected at a particular carcass decreases with the number of carcasses available at the time.

From the consumer point of view, a focal consumer is always in the basin of attraction of some resource; however, when resources are scarce it is unlikely that it will be close enough to detect the nearest resource. On the other hand, when resources are abundant, the area of the basin of attraction can be very small compared to the focal consumer's detection area, limiting the pool of potential consumers that might share the resource. In Figure. 4 and 5 we provide a comprehensive view of the dependence of a focal consumer's encounter rate *ε* on resource density. In Appendix A, we provide the details of a mathematical analysis of the model and rigorously demonstrate certain prominent features of the relationship: namely, the asymptotic power law in both the scarce and abundant resource regimes, as well as in the small and large distance of detection extremes. Furthermore, we provide an approximate formula for the resource density *𝓁*_*_ that leads to the maximum number of encounters for a given distance of detection and consumer density.

**Figure 4:**
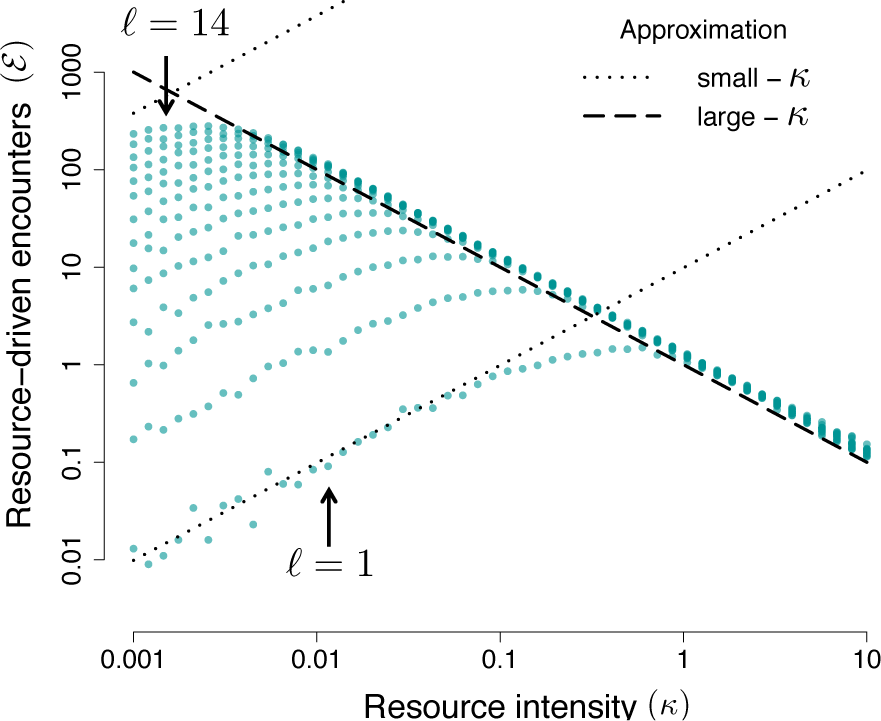
Encounter rate behavior. Each dot represents the average over 1000 simulations. The intersection of the dotted and dashed lines is the order-of-magnitude estimate described in the Results, Section 3.1.

**Figure 5:**
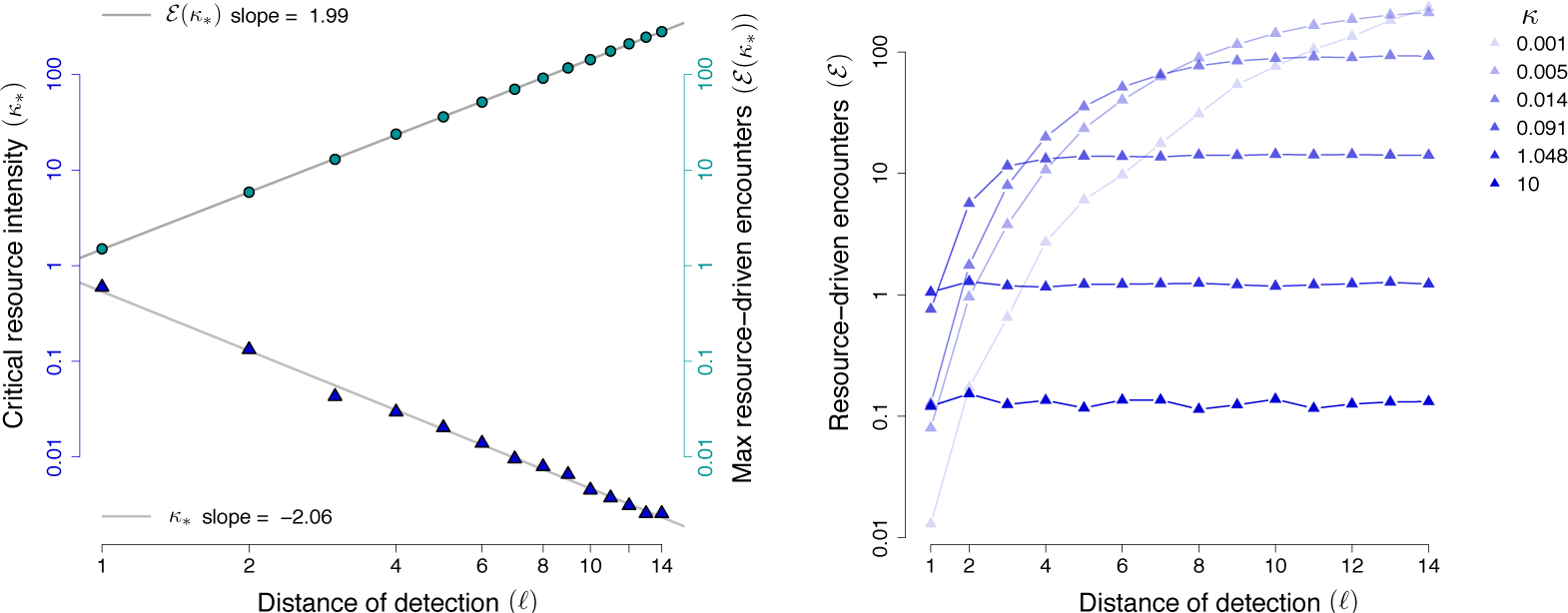
Left: In teal (circles), the maximum encounter rate as a function of *ℓ*. In blue (triangles), the resource intensity that yields the maximum. Right: Resource-driven encounters as a function of distance of detection. Lighter-to-darker shading corresponds to increasing values of ρ.

*Asymptotic results.* In Figure 4 we see that *ε*, the expected number of encounters for the focal consumer, has a very regular power law behavior in both the small- and large-*κ* regimes. Regardless of the value of *𝓁*, all of the encounter rate curves overlap in the large-*κ* regime. For small *κ*, the log-log slope is the same for all *ℓ*, but the leading coefficient differs. In Theorem A.1 we demonstrate that when resources are scarce (small *κ*), the resource-encounter function is asymptotically linear in *κ*. Furthermore we are able to establish the leading coefficient, yielding the *small-*κ* approximation ε ≈ ρκπ*^2^*ℓ*^4^ which is also validated by simulation. For example, in Figure 4, the lower black dotted line is the small-*κ* approximation when *ℓ* = 1 and we see good agreement for **κ* <* 0.1. When resources are abundant the analysis leads to an unsolved problem in spatial point process theory concerning the distribution of Voronoi cell (basin of attraction) sizes in tessellations generated by Poisson spatial processes. Nevertheless, we argue that ε scales with *κ*^−1^ in the abundant resource limit. Following the discussion in Section A.3, we present the *large-*κ* approximation ε* ~ *κ/κ* (Figure 4, black dashed line). The correct leading coefficient appears to be larger than *ρ*, but we were unable to obtain the exact value by mathematical analysis.

*Characterizing the encounter rate peak.* For reasons discussed in Section 3.4, perhaps the most important “landmark” of the resource-encounter function is its peak. Unfortunately it is difficult to directly analyze the magnitude of the peak and the corresponding critical resource density. However, there is a natural first-order estimate that involves the small- and large-κ approximations. Solving for their intersection yields the estimate *κ*_*\**_ ≈ (1/*π*)*ℓ*^−2^ and ε(*κ*_*\**_) ≈ *ρπℓ*^2^ where *κ*_*+*_ is the resource intensity that leads to the maximum resource-driven encounter rate. From Figure 4, it is clear that this is an overestimate, but not dramatically so. Using 1000 simulations at an array of *κ* and *ℓ* values, we found the following estimates using a linear regression:

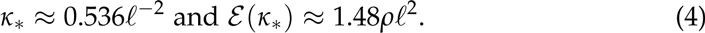

While the exponents align well with the intersection of the small- and large-*κ* approximations, we were unable to obtain a satisfactory explanation of the leading coefficients through direct mathematical analysis.

*Dependence on distance of detection.* In addition to characterizing the encounter rate's dependence on *κ*, we are also able to obtain an asymptotic understanding of the dependence of the resource-driven encounter rate on the maximum distance of detection parameter *ℓ*. As *ℓ* → 0, the encounter rate function behaves like ℓ^4^ (Theorem A.1). As might be expected, this function is monotonically increasing in *ℓ* and saturates to a limiting value for large *ℓ* (Theorem A.4). The corresponding simulation results are displayed in the right panel of Figure 5. The limiting value corresponds to the expected area of the basin of attraction in which the focal consumer resides. As explained before, this exact value is not known, but it scales like *κ*^−1^, which is why the limiting values in Figure 4 are largest for the smallest values of *κ*.

### 3.2 The relationship between resource density and site visitation

The mathematical model makes predictions about both full-population scale encounter rates and local single-resource site encounters. For the latter, from the perspective of a given carcass site, the model predicts that the maximum number of visitors should be observed when the resource density is the lowest. This is because in the sparse resource-density regime there is little to no competition for consumers. As the resource-density increases, the expected number of visitors should decrease. We consulted the ENP data set to investigate whether this effect can be observed for jackals and their tendency to visit carcasses that seasonally vary in abundance.

In the study area [15], the number of carcasses available for jackal scavenging varies seasonally (Figure 6, left panel inset). Between February and April, there is a resource pulse resulting from annual anthrax outbreaks in the local zebra population. These outbreaks occur during the end of the wet season [14, 16]. The timing and severity of anthrax outbreaks appears to be different between 2009 and 2010. The difference in severity provides an opportunity to make comparisons between the same months of the year but with very different numbers of available carcasses. In March and April, for example, the average number of jackals observed at carcasses decreases markedly from 2009 to 2010 when there are more carcasses available. In fact, in eight out of the eleven months where pairwise comparisons are possible, the average number of jackals observed at a carcass decreased when more carcasses were available in that month (right panel of Figure 6).

**Figure 6:**
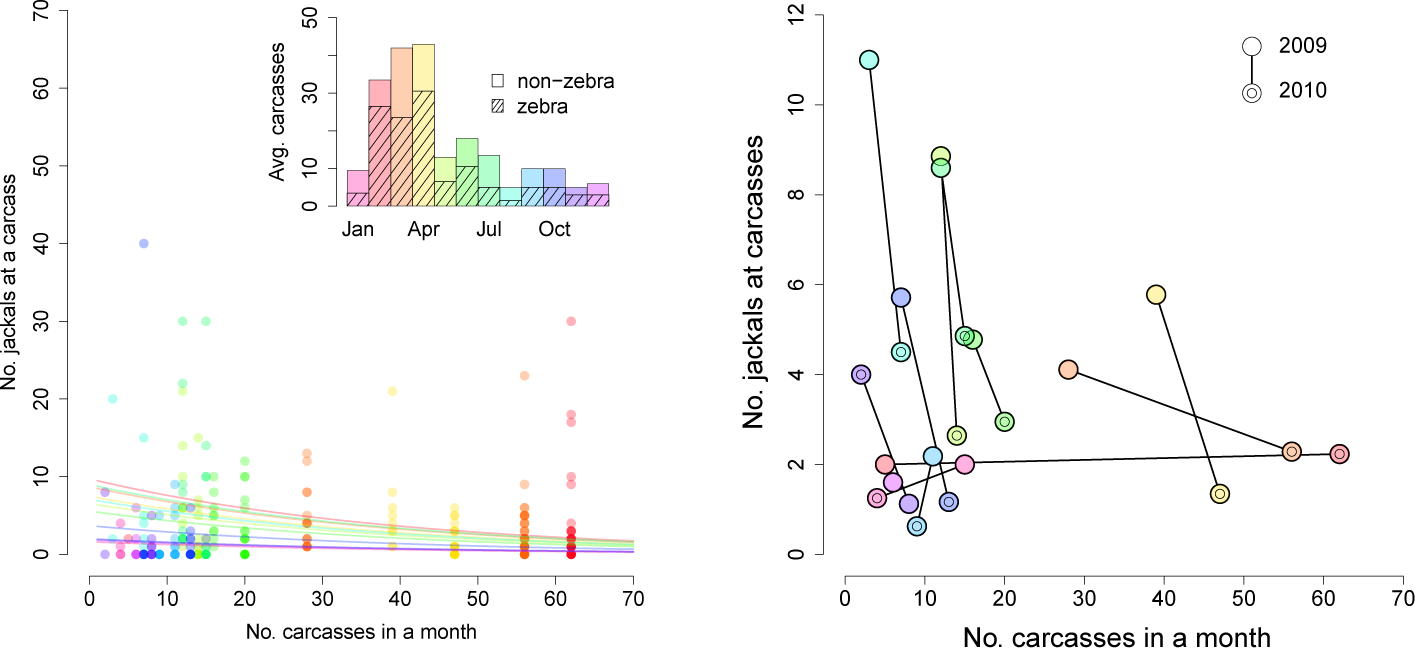
Monthly carcass availability and jackal visitation to carcass sites. Left: For each month-year pair, the number of observed carcasses is counted (x-axis). We plot the number of jackals recorded at each carcass versus the number of observed carcasses in the corresponding month. Jackal counts at carcasses are color-coded by month of the year as indicated by the bar chart in the upper right corner. Points are shaded so that darker shading indicates more observations. Regression lines are plotted for each month in the corresponding color. Inset: The monthly average number of observed total carcasses (solid bars) and zebra carcasses (striped bars). Right: For each month-year pair, the average number of jackals observed at carcasses is plotted. Repeated months are connected by lines.

To employ a more quantitative statistical test, we fit a Poisson general linear model with the number of observed carcasses as a predictor variable, and the number of jackals visiting a carcass as the response variable (available from January 2009 to November 2010). We also included predictor variables for each month of the year to allow for variation in environmental effects (e.g. wet/dry season), population processes (birth pulse, dispersal, etc.) and challenges in data collection that likely affect the expected number of jackals observed at carcasses. To be precise, let *y*_*i*_ be the response variable for the number of jackals observed at a carcass when there are *i* carcasses. Then

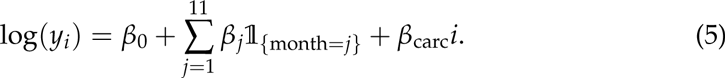

For example, in an April with *i* total carcasses, the expected number visitors observed at a carcass would be exp(*β*_0_ + *β*_3_ + *β*_carc_*i*). Using this statistical model, we found a significant negative correlation between the number of observed carcasses and the observed number of jackals at a carcass (*β*_carc_ = −0.025, 95% CI: [−0.029, −0.021]).

### 3.3 The relationship between defendable territory size and the distance of detection and response

Though there are three parameters in the mathematical model, we found that there are truly only two degrees of freedom in the parameter space. As shown in Appendix A.3, for every triplet (*ρ, *κ*, ℓ*), there is a corresponding triplet (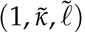), where 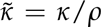 and 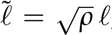, such that the expected numbers of encounters for the focal consumers are the same, i.e.

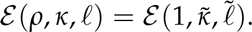

Notably, 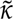 and 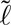 are nondimensional quantities and all formulas introduced in the previous section can be expressed using them:

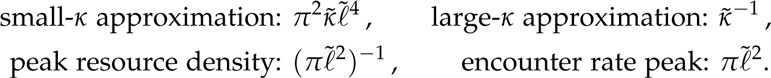

Both nondimensional quantities have informative biological interpretations. While it is straightforward to understand the significance of 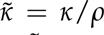 (the ratio of the resource density to the consumer density), the meaning of 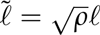 is more subtle. If we imagine dividing the landscape into even partitions, one for each consumer, then each consumer would be allocated a region of area 1/*ρ*. If the regions are square, then 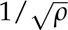 would be the length of each side and we can view 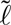 as 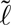 = 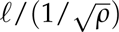 so that it is the ratio of the distance of detection to the typical length of a consumer's space allocation. In biological terms, we might think of these regions as the consumers' defendable territories and therefore 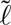 is roughly the number of defendable territories a consumer is willing to cross in order to visit a resource.

Our estimate of the resource density, *κ*, is based on carcass surveillance data from the Berkeley Etosha Anthrax Project during 2009 and 2010. The average number of carcasses recorded each month (left panel of Figure 6) was divided by four to get a weekly number of carcasses available. Since not all carcasses are observed, we followed Bellan et al. (2013) [26] in multiplying by a scaling factor of four to account for expected unobserved carcasses. We then divided the expected weekly number of carcasses available in each month by the area of the study region from Bellan et al. (2012) [15], roughly 1000 km^2^. This area contains all of the locations where carcasses were observed and jackal positions recorded. The resulting *κ* estimates ranged from 0.005 km^−2^ (August and November) to 0.043 km^−2^ (April).

As suggested by the nondimensionalisation argument above, we interpret *ρ* as the density of defendable jackal territories. Non-overlapping jackal territories in ENP were estimated to be between 4 km^2^ to 12 km^2^. This is comparable to estimates that were made for jackal populations in coastal Namibia (0.2 - 11.11 km^2^ [27]) and South Africa (3.4 - 21.5 km^2^ [28]). Noting the observation from Bellan et al. (2012) that jackals are “unusually dense” in ENP [15], we set the typical jackal territory size to be 5 km^2^, so that *ρ* = 0.2 km^−2^.

The interpretation of the parameter *ℓ* from the data requires some discussion. In the mathematical model, *ℓ* is the maximum distance at which a consumer can detect and then respond to a resource. We can think of the model as assuming that the probability of detecting a resource is one within a distance *ℓ* and zero outside that distance. Of course, in reality, this detection probability likely decreases steadily as a function of distance. Rather than identify a specific value that we definitively claim to be the best estimate of *ℓ*, we used the jackal movement data to find a range of reasonable values.

In Figure 7, we display a scatter plot of all jackal average positions relative to known carcasses and mark each with a teal dot or a gray x depending on whether the jackal visited the carcass or not. Jackals were observed to visit known carcass sites as far as 15 km away, but a large majority of carcasses visited were in a range of 0 to 4 km. As expected, the probability that a jackal visited a resource decreased with distance, but it is not known whether this was because the jackals were not aware of more distant carcasses, or because there were other carcasses or alternate resources nearer by. In Section 3.4 we use the two values *ℓ* = 4 and 10, and the associated encounter-rate curves are displayed in Figure 8. Each was generated by averaging the results of 10,000 simulations at each of 300 values for *κ*.

**Figure 7:**
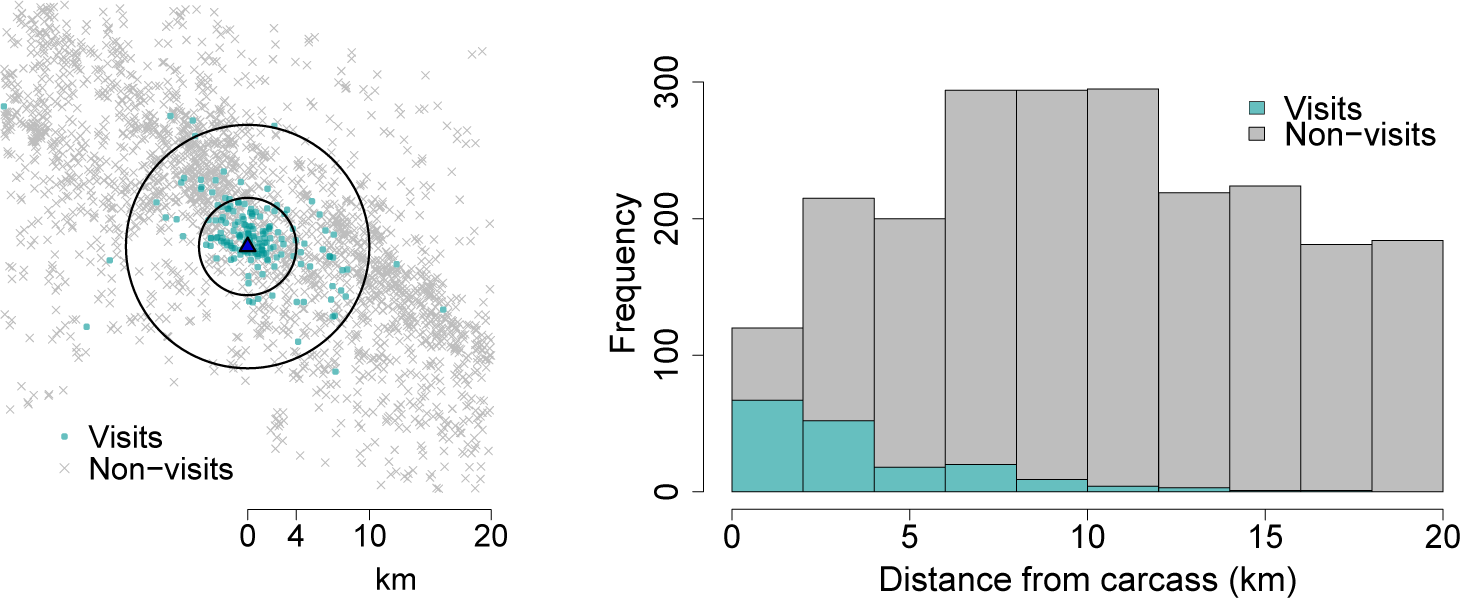
**Left:** Relative positions of jackals plotted with respect to the location of each known carcass. Jackal locations were calculated as the average of their GPS pings that occurred between two days before and after the carcass was estimated to be present. **Right:** Stacked histogram for the times that jackals chose to visit a carcass (teal bars) and the times that jackals refrained from visiting a carcass (gray bars).

### 3.4 Placing model results in the context of Disease Ecology

In Section 2.2, we described our stochastic small-population model for pathogen invasion. We say that an invasion is “successful” if it achieves a population level equivalent to what would be the endemic equilibrium of the deterministic version of the model. There exists an explicit formula for the probability of invasion, but it is difficult to interpret in terms of the parameters of the model. So, following Ball & Donnelly [24], we use an approximation for the true value (see Equation 3 and further discussion in Appendix A). This reduces our analysis to determining whether the total rate of transmission (which is affected by the resource-driven encounter rate) is greater than the disease-related mortality rate *v*.

To assess whether a change in the consumer encounter rate is “large” in the context of jackals and rabies, we followed Rhodes et al. [29] in establishing a background rate of pathogen transmission (b = 1 wk^−1^) and a disease-related mortality rate (*v* = 1.4 wk^−1^) yielding the reproductive ratio 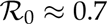 Since 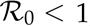, rabies is found to be sub-critical. To connect the resource-driven encounter rate at a given time *t*, ε(*t*), to the pathogen-transmission model, we first note that not all encounters involving infectious and susceptible individuals lead to a new infection. For example, in our model, a resource-driven encounter is defined to occur if two individuals visit the same resource site in the same week, but this does not mean they visit concurrently. Even if they visit concurrently this does not ensure pathogen transmission. Define *p*inf to be the probability that a resource-driven encounter results in transmission. Then our expected number of new infections arising from a single infectious individual is *b* + *p*_inf_^ε*(t)*^ and the reproductive ratio is

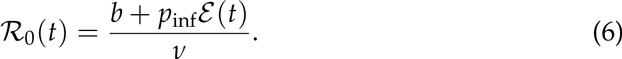

Using the month-by-month encounter rate values appearing in Figure 8, we calculated the time-dependent reproductive ratio for six scenarios and displayed them in Figure 9. The left and right panels correspond to the distance of detection choices *ℓ* = 4 and *ℓ* = 10, respectively. In each case, we varied the probability of infection parameter *κ*_inf_ to demonstrate its impact on the final result.

**Figure 8:**
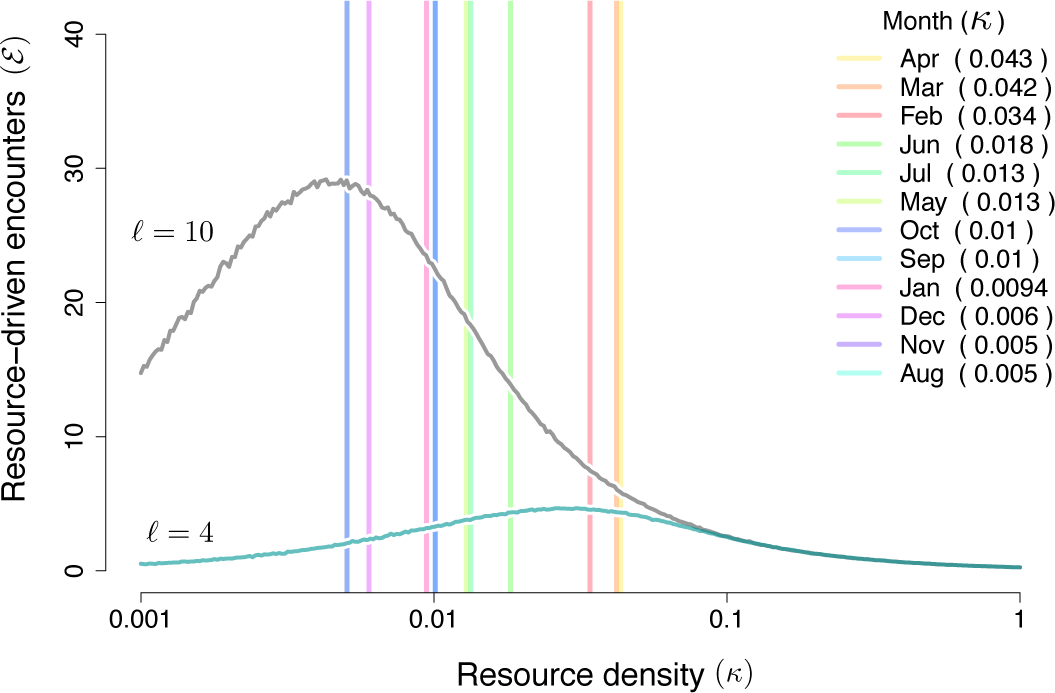
Simulated number of resource-driven encounters for two choices of the the detection distance parameter over a range of resource intensities. The vertical lines indicate the estimated month-by-month carcass densities observed in the ENP data set.

**Figure 9:**
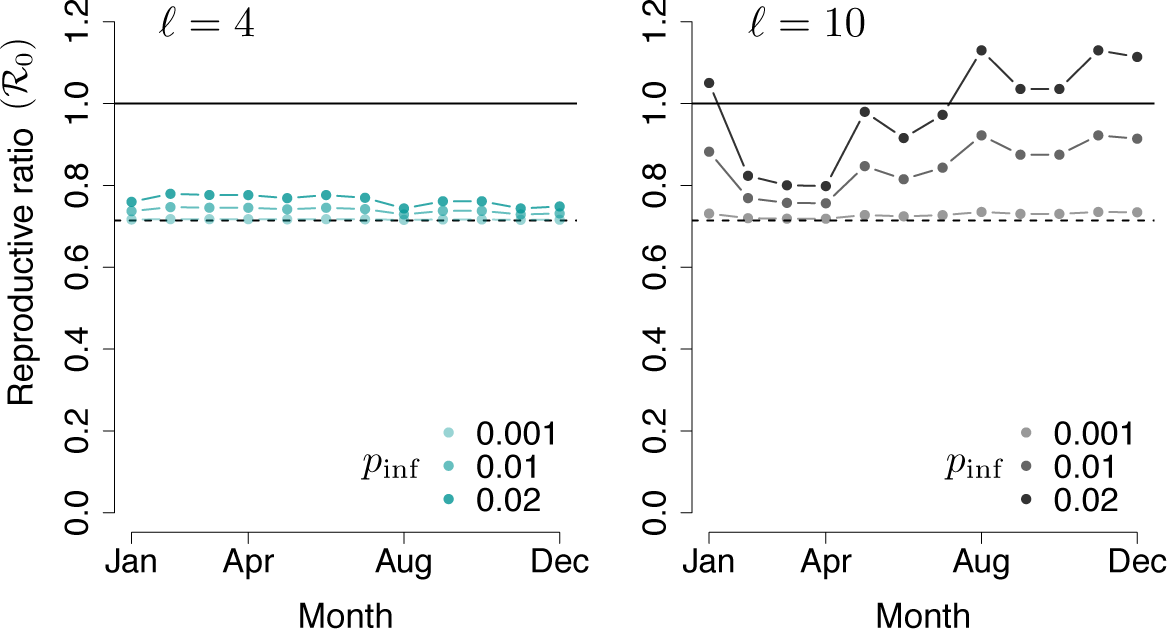
Time-dependent reproductive ratio based on the corresponding number of resource-driven encounters for each month in Figure 8 with *b* = 1, *v* = 1.4 and *ℓ* = 4 (**Left**) or *ℓ* = 10 (**Right**).

**Table 1:** Parameters used in the Disease Ecology analysis.

|  | Value | Units | Definition | Source |
| --- | --- | --- | --- | --- |
| $b$ | 1 | wk<sup>-1</sup> | transmission rate | [29] |
| $\nu$ | 1.4 | wk<sup>-1</sup> | rabies mortality rate | [29] |
| $\kappa$ | 0.005 - 0.043 | km<sup>-2</sup> | carcass density | ENP data analysis |
| $\rho$ | 0.2 | km<sup>-2</sup> | jackal territory density | [15, 28, 27] |
| $\ell$ | 4-10 | km | max distance of detection | ENP data analysis |
| $\tau$ | 1 | wk | timescale of resource availability and visitation | ENP data analysis |

When *ℓ* = 4, the resource density for each month is below the critical resource density *κ*_*_, i.e. the density for which the maximal encounter rate occurs. So an increase in resource density leads to increases in the resource-driven encounter rate and resulting reproductive ratio, regardless of the *p*_inf_ value. However, because the peak of the encounter rate curve is relatively low (approximately five per week, see Figure 8) the reproductive ratio remains below the critical value of one. On the other hand, when *ℓ* = 10, most of the monthly resource densities are greater than *κ*_*_. In those cases, increases in resource density lead to decreases in the resource-driven encounter rate and resulting reproductive ratio. In this regime, we see that the months with low carcass availability are most vulnerable to pathogen invasion.

We note that the magnitude of change in 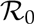 is directly dependent on the estimate for jackal territory density *ρ* (recall Equations 4). If *p*_inf_ = 0.02 and if *ρ* = 1 instead of *ρ* = 0.2, for example, then the April 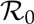 for *ℓ* = 4 would be approximately 1.07. This constitutes a setting where indirect induction of disease is possible. A similar modification for *ℓ* = 10 would result in an August 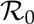 of 2.86. These results can readily be translated to a probability of successful invasion over the course of a resource increase of duration *T*. As described in Section 2.2, successful pathogen invasions arrive according to a Poisson process with rate γ_spillover_p_invasion_. Assuming the transmission rate is constant over the period of interest, the probability of invasion is 1 - exp (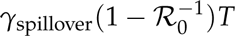).

## 4 Discussion

In this work, we have developed a framework for analyzing the impact of changes in resource availability on the rate of conspecific encounters among consumers and express our results in the context of disease ecology. Given a landscape of consumers and resources we essentially ask the question: would adding one more resource site lead to more or fewer encounters among the consumers?

We have proposed a novel consumer-resource interaction model to investigate this question. Through a combination of numerical simulation and mathematical analysis, we have identified and characterized two qualitatively distinct parameter regimes. In a scarce resource regime, adding more resources leads to more consumer-consumer encounters; in an abundant resource regime, adding more resources leads to fewer consumer-consumer encounters. The utility of our model is that it can be used to predict the qualitative dynamics of a system once certain fundamental parameters are estimated: the consumer density (*ρ*), the resource density (*κ*) and the maximum distance of detection and response (*κ*).

To work through a specific case study, we used location data for a population of jackals and the carcasses upon which they scavenge in Etosha National Park. While some model parameters (*κ* and *ρ*) are fairly straightforward to estimate, others are not (see Section 3.3 for our approach to estimating the parameter *ℓ* in particular). One notable challenge that arises is that the definition of an “encounter” is intrinsically subjective, depending strongly on the question of interest (also see Gurarie et al. [2] for a full discussion on this point). To relate our resource-driven encounter rate to a rate of pathogen transmission from infectious to susceptible individuals, we introduced a corrective “probability of infection” factor p_in_f. Because pathogen transmission is essentially impossible to directly observe, proper inference for such a parameter would likely require population-level disease incidence data that does not currently exist. In response to this uncertainty in parameter values, we display model results that emerge from a range of reasonable values for both *p*_inf_ and *ℓ*. The key takeaway is that for certain combinations of biologically relevant parameters, we confirm that small changes in the resource landscape can lead to substantial changes in pathogen transmission dynamics. In fact, we show that sudden scarcity of a resource can have a larger effect on encounter rates than a resource pulse.

Building upon existing investigations into how changes in resource and consumer densities induce changes in disease dynamics, our work suggests that the relationship between territory size and the distance of resource detection plays a crucial role in determining infectious disease outcomes. To use the present context for an example, we note that jackals may use visual cues from vultures to identify carcass sites ([30] and anecdotal observations by an author and colleagues). If vulture populations decline, as has now been documented in both Asia and Africa [31], the detection distance for jackals could decrease, potentially causing a pathogen invasion regime shift. Interestingly though, the specific example of declining vulture populations exemplifies the complexity of consumer-resource interactions. In an experiment conducted by Ogada et al. [32], the authors found that there were increased encounters among mammalian scavengers when vultures could not see and react to carcasses (in contrast to the right hand panel of our Figure 5).

### 4.1 Opportunities for integrating more detailed animal behavior

The complex relationship between resource allocation, consumer behavior, and pathogen spread deserves further study. We constructed our model to be detailed enough to examine our primary question, but simple enough to permit rigorous mathematical analysis. While there are many ways to extend the model to account for more nuanced behavior, we highlight a few.

*Resource detection and selection.* There are other natural models for the consumer's ability to detect resources, as well as for the algorithm determining which resource is visited, if any. For example, one could posit that there is imperfect detection and that the probability of detection decreases with a consumer's distance from the resource. Also, one could relax the restriction that the consumer always picks the closest detected resource. An informal investigation suggested that, as long as we pose assumptions consistent with those outlined at the beginning of Section 2.1, adopting alternative model specifications does not change the qualitative description of our results reported in Section 3.1. We opted for the version that yields the most explicit analytical results, but note that changes to model assumptions would likely change the value of the critical resource density *κ*_*\**_ as well as the height of the associated encounter rate peak.

One major factor that we did not consider is heterogeneity in the resource sites. Variation in resource quality, geographical characteristics and local environmental factors can affect the model through multiple parameters. Resource sites that attract vultures might be detectable from larger distances than resources that do not, causing variation in *ℓ*. Small carcasses may be rapidly depleted, decreasing *<*_*1*_, and may not satiate consumers, decreasing *<*_*2*_. In terms of the selection algorithm, a consumer might not choose the closest available resource if one of greater quality is just a little bit farther away. Our model assumes uniformity in resources and, compared to predictions that would follow from each of these possible modifications, it produces a lower variance in the number of visitors to a given site.

The reduction in variance is significant in the following sense. As we report in Appendix A, Equation 9, the number of visitors to a given site is Poisson distributed with a mean parameter that decreases with the total number of available resources. While this prediction for the mean is consistent with the available carcass visitation data (see Figure 6), the variance of a Poisson random variable is much less than the variance of the observed distribution. In February of 2010, for example, there are values as high as thirty when the mean is less than five. However, this is extremely unlikely for a Poisson distribution. To be precise, if 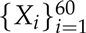 are independent and identically distributed Pois(5) random variables, then *P(max_i_X_i_* > 20) < 0.0001.

*Modified behavior of infectious individuals.* Behavioral changes associated with the disease status of an individual may affect its expected encounter rate. Developing a theory for susceptible-infectious encounter rates that considers both types of individuals will be especially important for infections that alter host behavior (e.g. rabies). Specifically, we note that the manner in which a rabid animal detects and selects resource sites could be much different than that of a susceptible individual.

*Off-site encounters.* At present, our model considers the relationship between resource availability and the consumer encounter rate specifically at resource sites. However, a change in resource availability will likely influence other types of encounters as well. For example, when consumers are forced to make long treks to scarce resources, they may be exposed to unfamiliar individuals. Distinguishing between typical encounters (e.g. with family members and territorial neighbors) and unique encounters with new individuals could be important for determining transmission dynamics [33, 13].

*Dynamic population counts.* We considered a fixed population density (i.e. ρ, the jackal territory density). However, population sizes change on multiple time scales. Jackals have birth pulses that will change the local jackal population size on an annual basis (although, pups may not contribute dramatically to pathogen spread). In the long term, consumer population size may respond to resource availability; when resources are abundant more consumers can be supported in the same area. This allows for smaller territories (increases in *ρ*). Specifically in Etosha, zebras are attracted to a grassland foraging area south of the salt pan. The jackal density in this area may be higher than the density in other areas due to greater average resource availability. Considering Figure 4 and Equations 4, we see that if the consumer density varies with the resource density, then there are two competing effects: while increasing *κ* can decrease the number of encounters for fixed *ρ*, a simultaneously increasing *ρ* can overcome this effect.

### 4.2 Using seasonality as a tool for investigation

Large-scale ecological experiments are expensive and challenging to conduct. It can therefore be very useful to observe and characterize systems with naturally changing resource conditions [9, 11]. Being able to observe the same system in multiple states provides the opportunity to investigate responses to the altered system components while keeping other characteristics constant. Seasonality, in particular, is frequently observed in time-series data for incidence of disease and has been shown to affect infection rates through multiple mechanisms. Seasonal changes can result in a fluctuating population size and affect both the quantity and type of conspecific interactions. Moreover, periodic changes in population count due to birth pulses [34] and migration [35] have both been shown to affect the potential for infectious disease outbreaks. We believe that coupling temporally varying environmental information in multi-state systems with rigorous analysis of GPS location data can provide a basis for more mechanistic models of consumer response to resource change. Ultimately this can lead to more meaningful and more accurate predictions for the consequences of habitat alteration.

## Acknowledgements

The authors would like to thank Wayne M. Getz, Benjamin M. Bolker, Craig W. Osenberg, and Andrew M. Hein for their support and thoughtful conversations in the development of this work.

## Funding Statement

This work was supported by the International Clinics on Infectious Disease Dynamics and Data (ICI3D) program, with funding from the South African Centre for Epidemiological Modelling and Analysis (SACEMA), the African Institute for Mathematical Sciences (AIMS), and the US National Institutes of Health (NIGMS award R25GM102149 to JRCP and A Welte). Data collection for this work was conducted by the Berkeley Etosha Anthrax Project with support from an NSF-NIH Ecology of Infectious Disease Grant (award GM83863 to WM Getz) and with assistance from the Etosha Ecological Institute, (Namibian Ministry of Environment and Tourism). Additional funding was provided from the Center for Inference and Dynamics of Infectious Diseases (MIDAS-NIGMS, award U54GM111274 to ME Halloran). JRCP was supported by the Research and Policy on Infectious Disease Dynamics (RAPIDD) Program of the Fogarty International Center, National Institutes of Health and Science and Technology Directorate, Department of Homeland Security. SEB was supported by NSF-NIH (GM83863 to WMG), MIDAS-NIGMS (award U01GM087719 to LA Meyers and AP Galvani) and NIAID (K01AI125830 to SEB). SAM and JMF were partially supported by the Army Research Office (ARO 64430-MA). RKB was partially supported by the National Science Foundation under Grant 0801544 in the Quantitative Spatial Ecology, Evolution and Environment Program at the University of Florida, and the CLAS Dissertation Fellowship, funded by the Kenneth and Janet Keene Endowed Fellowship.

The authors declare that they have no competing interests.

## Author Contributions

SEB collected much of the data and asked the motivating question for this research. SAM, RKB and JMF collaborated in the mathematical model development. Numerical simulations were written and executed by RKB and JMF. Mathematical analysis was performed by RKB and SAM. Statistical analyses were conceived and performed by all five authors. RKB, JRCP, SEB, and SAM developed the framing of the manuscript, description of biological intuition, and presentation of results. The manuscript was written by RKB and SAM with substantial input and edits from SEB, JRCB and JMF.

## A Mathematical Analysis

The simplicity of the resource-driven encounter model invites a rigorous asymptotic analysis. More than demonstrating the non-monotone relationship between resource density and the consequent encounter rate in the consumer population, we can obtain the exponents of the power laws that govern the relationship.

In what follows (and in the main text) when we write *ϕ(x)~ x*^*a*^ as *x → a*, we mean that there exists some constant *C* ∊ (0, ∞) such that

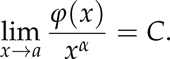

For example, a result we will use below is that if *γ* ~ Pois(λ) for some λ > 0, then 𝕡 { *γ* > 1} ~ λ 2 as λ → 0. This is because

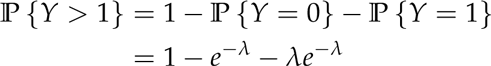

and using the Taylor series expansion for the exponential (or simply L’Hôpital’s rule), we have

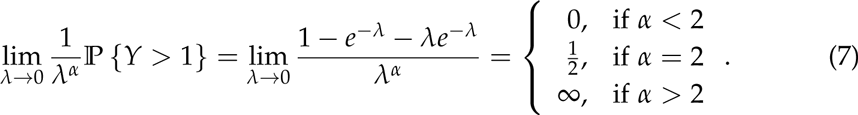

For higher order terms we will use Big-Oh notation: we say that *f(x)* = *O*(*g*(*x*)) near *x* = *a* if there exist constants *C >* 0 and *L >* 0 such that if |*x – a*| *< L*, then |*f(x)*| ≤ *C*|*g(x)*|.

As in the main text, *κ* and *ℓ* denote the resource intensity and maximum distance of detection respectively. In the presentation of our results we will assume that the consumer density *ρ* = 1. In Section A.3 we will discuss how to modify the results when **ρ* ≠* 1.

We take the domain 𝒪 to be a circle of radius *R >* 3*ℓ* centered at the origin. There is a *focal consumer* located exactly at the origin. Resources are distributed throughout 𝒪 as a Poisson spatial process with intensity *κ*. Other, non-focal consumers are distributed throughout 𝒪 as a Poisson spatial process with intensity one. Let 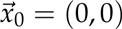 = (0,0) and enumerate the non-focal consumer locations 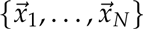 where *N* ~ Pois(|𝒪|). Furthermore let 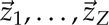 be the resource locations where *Z* ~ Pois(κ|𝒪|). For each pair 1 < *i < N* and 1 < *j < Z*, let 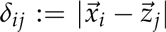. For each i ε {0,…,N}, let η_i_:= {j: δ_ij_ = min_1≤j≤z_δ_ij_}. In other words, *η*_*i*_ is the index of the resource that is closest to the *i*th consumer. For notational expediency we will write the index of the resource closest to the focal consumer, *η*_*0*_, to simply be *η*.

In the above notation, we can express **β**, the number of resource-driven encounters experienced by the focal consumer, to be

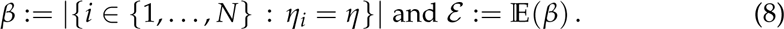

Given a set of resource locations, it is useful to think of the landscape partitioned according to the associated Voronoi tessallation. That is, neglecting a set of measure zero,

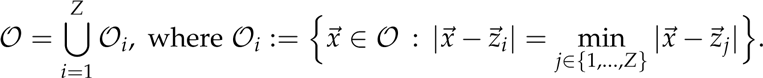

We say that 𝒪_i_ is a *basin of attraction* for resource *i*: all consumers located in 𝒪_i_ will choose resource *i* as their resource to visit if it is within their detection radius. We define 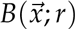 to be the circle of radius *r* centered at the location 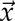. Then the distribution of the encounter variable **β** conditioned on a given resource landscape 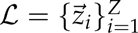

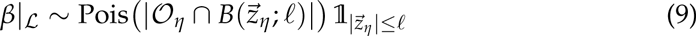

where we recall that *η* is the index of the resource chosen by the focal consumer and ퟙ_*A*_ if the event *A* occurs, and is zero otherwise.

### A.1 Small resource density and/or small detection distance

#### Theorem A.1.

*Let ε = ε(κ, ℓ) be defined as in (8). Then ε ~ *κ* and ε ~ ℓ*^*4*^ *as *κ* and *ℓ* go to zero, respectively. To be precise*,

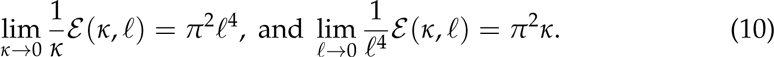

*Proof.* We first introduce some notation. Let *N(r)* and *Z(r)* denote the number of consumers and resources within a distance *r* of the focal consumer. We proceed by conditioning on the number of resources that are near the focal consumer. We partition the sample space Ω as follows:

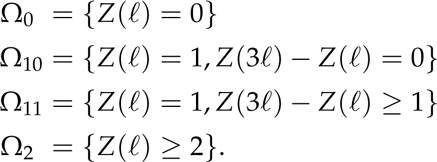

Naturally, it follows that 𝔼(*β*) = Σ*i* 𝔼(*β*|Ω*i*) 𝕡 {Ω*i*} and we will find that the dominant term is the one associated with Ω_10_. Looking at the other terms, first observe that 𝔼(*β*|Ω_0_) = 0 since, if there are no resources to consume, the focal consumer will not have any encounters.

To deal with the event Ω_2_, we apply Eq 7 above, noting that the number of resources within detection distance of the focal consumer has the distribution Z(*ℓ*) ~ Pois (*κπℓ*^2^). It follows that for small *κ*, 𝕡{Ω_2_} ~ *κ*^2^ and for small *ℓ*, 𝕡{Ω_2_} ~ ℓ^4^. To bound the conditional expectation 𝔼(*β*|Ω_2_), observe that the number of resource-driven encounters experienced by the focal consumer must be less than or equal to the number of consumers that are located within a distance of *ℓ* of the focal resource (the resource chosen by the focal consumer). Because this is a region of size *πℓ*^2^, we have 𝔼(*β*|*Ω*_2_) ≤ *πℓ*^2^. Together we have that

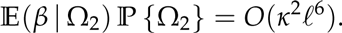

For the event Ω_11_ we again exploit that, when the detection distance or resource density is small, it is unlikely that there will be more than one resource near the focal consumer. By independence of the resource distribution in disjoint regions

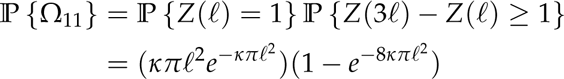

since the area or the annulus covering the region that is between a distance or *ℓ* and 3*ℓ* of the origin is 8*πℓ*^2^. It follows that 𝕡{Ω_11_} = *O*(*κ^2^*ℓ*^4^*). Meanwhile, using the same upper bound on the number of resource-driven encounters experienced by the focal consumer, we have 𝔼(**β** | Ω_11_) *≤ πℓ*^2^. Therefore

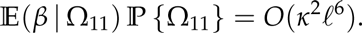

Turning our attention to the event Ω_10_, if there is only one resource in the cal consumer's detection radius, and the resource is the only one in the larger 3*ℓ* radius circle centered at the origin, then all consumers within a radius *ℓ* of the focal resource will choose the same resource as the focal consumer. In other words, the number of encounters conditioned on Ω_10_ is **β** | Ω_10_ ~ Pois(*πℓ*^2^). What was an upper bound in previous cases is now equality. It follows that 𝔼(**β** | Ω_10_) = *πℓ*^2^. To compute the event's probability we argue as before,

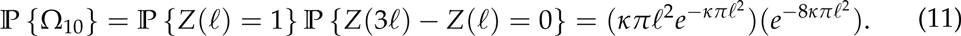

Therefore

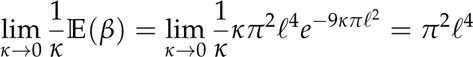

and

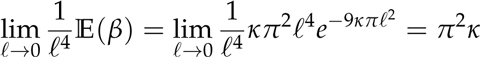

as claimed. □

### A.2 Analysis in the high resource and large distance of detection regimes

In the high resource density and large distance of detection we are unable to get exact results. This is due to a fundamental barrier in the analysis that we will describe below. In the high density regime we can provide what appears to be a lower bound on ε that, from the numerics, seems to scale with *ε* as **κ* → ∞*.

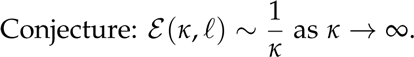

Our conjecture is based on the following heuristic. Recall that *𝒪*_*η*_ is the basin: attraction that contains the focal consumer. Then:

- Conditioned on the landscape of resources, the number of encounters experienced by the focal consumer is Poisson distributed with mean equal to its containing basin of attraction. Therefore 𝔼(**β**) = 𝔼(|𝒪_*η*_|).
- *Unconfirmed estimate:* 𝔼(|𝒪_*η*_||Z = *z*) ≥ |𝒪|/*z*.
- 𝔼(|𝒪|𝟙_z>0_/Z) ~ 1/*κ* as *κ* → ∞.

The third part of the heuristic is established by Lemma A.3 below. The second art of the heuristic is justified by the following.

#### Lemma A.2.

*Let a resource landscape be given as described above and let the total region* ^1^ *be partitioned according to a Voronoi diagram generated using the resource locations* 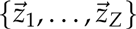. *We denote the areas of each of these basins of attraction* {*A*_1_*,…,A*_*Z*_}.

*Let 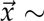 Unif (𝒪) be a random location in the landscape and define η to be the index 'the basin of attraction that contains this point. Then*

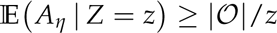

Remark 1. *Unfortunately, at this time, we are not able to extend the result to establish the claim that the basin of attraction specifically containing the origin has an expected area that is larger than 𝒪/z. Numerics strongly support this conclusion.*

*Proof.*

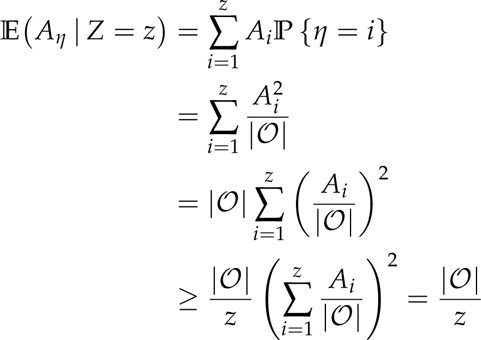

where, in the last line, we have used the Cauchy-Schwarz inequality. □

#### Lemma A.3.

*Suppose γ ~ Pois(ℓ). Then*

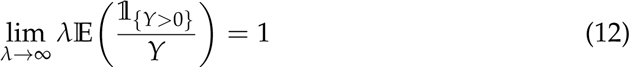

*Proof.* Recall the exponential integral function

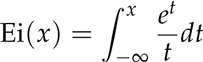

for *x* > 0, where the integral is taken in the sense of the Cauchy principal value. The exponential integral function can be written in terms of the series [36].

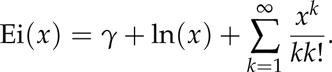

For large x, Ei(*x*) has the asymptotic expansion [37],

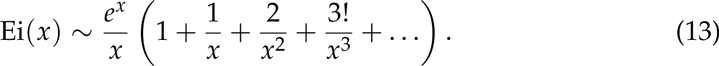

Following the suggestion of Grab and Savage [38], we note that

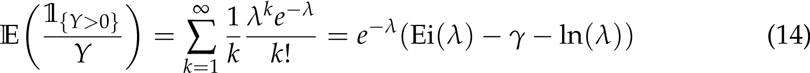

where *γ* is the Euler-Mascheroni constant. Combining (13) and (14) we arrive at the desired result. □

There is a fundamental mathematical barrier to making more progress on this problem. The preceding analysis reduces the problem to analyzing the distribution of areas of cells generated by Poisson Voronoi Tessellations, but this an outstanding mathematical problem [17]. In particular, we there is no known expression for 𝔼(|𝒪_η_|), the expected area of the basin of attraction that contains the focal consumer. This prevents us from obtaining a result for the high distance-of-detection regime that is explicit in *κ*.

#### Theorem A.4.

*ε*(*κ, ℓ*) is an increasing function in *ℓ* and

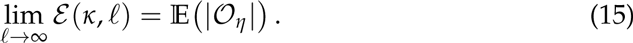

*Proof.* For a given *κ* > 0, let 𝑸_*κ*_ denote a landscape of resources generated by a spatial Poisson process with intensity *κ*. For each such landscape, let **β**|_*𝑸κ*_(*ℓ*) be the number of encounters for the focal consumer. As noted above in Equation (9), this is the number of consumers located in a radius *ℓ* of the focal resource multiplied by one or zero depending on whether the focal resource is within radius *ℓ* of the focal consumer. For a fixed landscape, note that

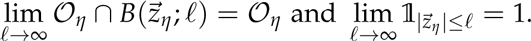

As such,

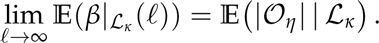

Because this holds for all 𝑸_*κ*_, the proposition follows. □

### A.3 Converting results for non-unit consumer density

All of the preceding results have been expressed under the consumer density assumption *ρ* = 1. Similarly all simulations were conducted with *ρ* = 1. Extending the earlier notation, let *ε* (*ρ, *κ*, ℓ*) be the expected number of encounters for the focal consumer for the given triplet of parameters. We claim that, although there are three fundamental parameters in the model, there are only two degrees of freedom in the parameter space. That is to say, given a triplet (*ρ, *κ*, ℓ*) there exists a unique pair (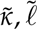) such that 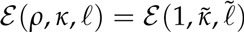. Namely,

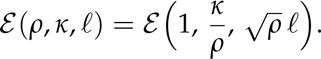

To see this, let *N*(ᴈ*ℓ*) and 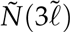 denote the number of consumers in the model for the parameter triplets (*ρ, *κ*, ℓ*) and (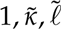) respectively. Let Z(ᴈ*ℓ*) and 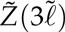 denote the same for resources. Because consumers and resources are distributed as Poisson spatial processes, these values completely define the system. Furthermore, because Poisson random variables are completely parameterized by their means, it follows that 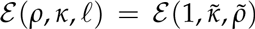 if 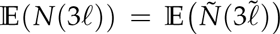 and 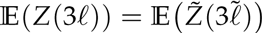. For the first constraint,

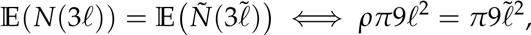

from which it follows that 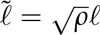. Meanwhile

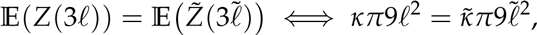

meaning that 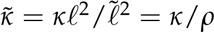

We note that this reduction of the problem to two parameters amounts to a nondimensionalization of the three parameter model. The units of *ρ* and *κ* are both [length]^−2^, while *ℓ* has units of [length]. As a result, 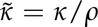 and 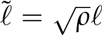 are both dimensionless.

Revisiting the theorems of the previous sections we have the results:

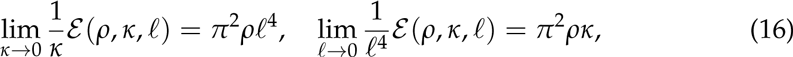

and

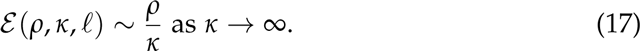

### A.4 Branching process approximation

There exist exact solutions to the hitting probability problem introduced in Section 2.2, however, such a presentation makes it difficult to understand how *κ*_invasion_ depends on *b* and *v*. Under the assumption that the size of the susceptible pool is very large with respect to the initial infectious population, it is common to introduce the approximation that (*N – I*) */N* ≈ 1 [39]. The modified rate functions for the CTMC are linear and take the form

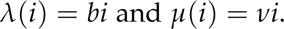

The infectious population process is then a Galton-Watson branching process. he only two outcomes for such a process are extinction or explosion to infinty. The analysis reduces to recasting the CTMC as a discrete time generation-by-generation branching process that is defined in terms of the *offspring distribution*, i.e., the distribution of the number of offspring an individual might have before lying. In our case, the “offspring” are the infections spawned by a single inividual. Since the infection events occur according to a Poisson process with rate parameter *b* and the death of the individual occurs at rate *v*, the number of successful infections before death is Geometrically distributed with success probability *m := b/*(*b + v*). One can then show that, under the assumption that *b* > *v*,

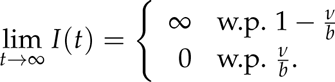

If *b ≤ v*, the process goes extinct with probability one.

